# Benchmarking freely available human leukocyte antigen typing algorithms across varying genes, coverages and typing resolutions

**DOI:** 10.1101/2022.06.28.497888

**Authors:** Nikolas Hallberg Thuesen, Michael Shantz Klausen, Shyam Gopalakrishnan, Thomas Trolle, Gabriel Renaud

**Affiliations:** Evaxion Biotech, Copenhagen, Denmark; Department of Health Technology, Section for Bioinformatics, Technical University of Denmark, Kgs. Lyngby, Denmark; Section for Hologenomics, Department of Biology, University of Copenhagen, Copenhagen, Denmark

**Keywords:** Human Leukocyte Antigen, next-generation sequencing, whole exome sequencing, typing resolution, algorithm, benchmark, depth of coverage, ancient DNA

## Abstract

The human leukocyte antigen (HLA) system is a group of genes coding for proteins that are central to the adaptive immune system and identifying the specific HLA allele combination of a patient is relevant in organ donation, risk assessment of autoimmune and infectious diseases and cancer immunotherapy. However, due to the high genetic polymorphism in this region, HLA typing requires specialized methods.

We investigated the performance of five next-generation-sequencing (NGS) based HLA typing tools with a non-restricted license namely HLA*LA, Optitype, HISAT-genotype, Kourami and STC-Seq. This evaluation was done for the five HLA loci, HLA-A, -B, -C, -DRB1 and -DQB1 using whole-exome sequencing (WES) samples from 829 individuals. The robustness of the tools to lower coverage was evaluated by subsampling and HLA typing 230 WES samples at coverages ranging from 1X to 100X. The typing accuracy was measured across four typing resolutions. Among these, we present two clinically-relevant typing resolutions, which specifically focus on the peptide binding region.

On average, across the five HLA genes, HLA*LA was found to have the highest typing accuracy. For the individual genes, HLA-A, -B and -C, Optitype’s typing accuracy was highest and HLA*LA had the highest typing accuracy for HLA-DRB1 and -DQB1.

The tools’ robustness to lower coverage data varied widely and further depended on the specific HLA locus. For all class I loci, Optitype had a typing accuracy above 95% (according to the modification of the amino acids in the functionally relevant portion of the protein) at 50X, but increasing the depth of coverage beyond even 100X could still improve the typing accuracy of HISAT-genotype, Kourami, and STC-seq across all five HLA genes as well as HLA*LA’s typing accuracy for HLA-DQB1. HLA typing is also used in studies of ancient DNA (aDNA), which often is based on lower quality sequencing data. Interestingly, we found that Optitype’s typing accuracy is not notably impaired by short read length or by DNA damage, which is typical of aDNA, as long as the depth of coverage is sufficiently high.

## Introduction

The human leukocyte antigen (HLA) region is a gene cluster of 45 genes located on chromosome 6, which encodes membrane bound proteins involved with antigen recognition in the human adaptive immune system. HLA genes can be divided into subclasses based on their structure and function. Generally, HLA class I proteins are found on the surface of most somatic cells and present peptides, originating from proteins produced within the cell, to CD8^+^ cytotoxic T lymphocytes (CTLs), while HLA class II proteins are found on antigen-presenting cells (APCs) and present exogenous peptides to CD4^+^ helper T-cells (see figure 1 **(A)**) (1, 2).

**Fig. 1.**
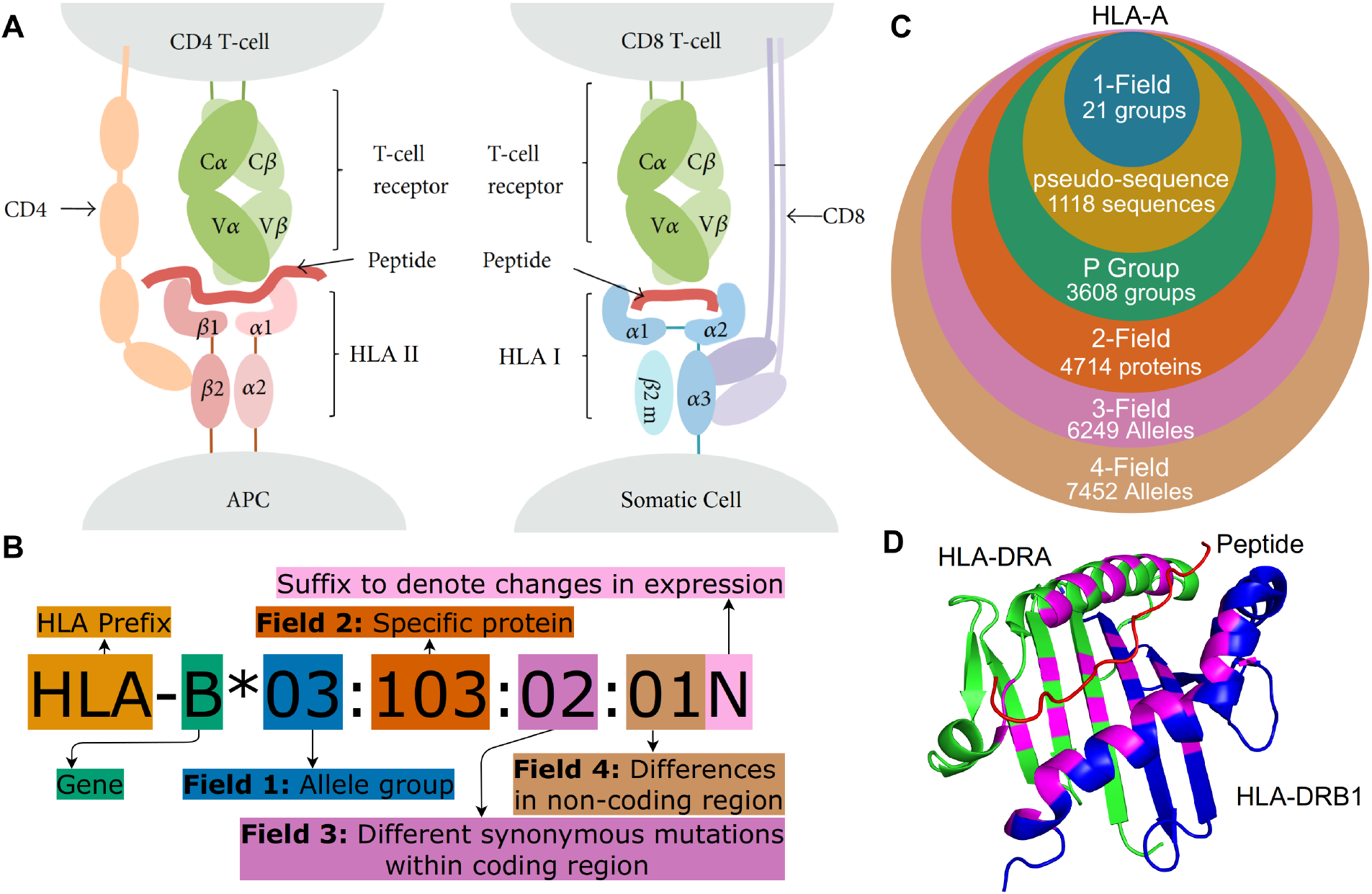
**(A)** Peptide presentation on APCs to (I) a CD4^+^ helper T-cell via an HLA class II protein and (II) to a CTL (CD8^+^ T cell) via an HLA class I protein. Both class I and class II molecules are heterodimers but for class II, the ARD is made up of one domain from each monomer and therefore encoded by two different genes. Figure adapted from (9). **(B)** HLA nomenclature shown with full four field (8-digit) resolution. Adapted from http://hla.alleles.org/nomenclature/naming.html. **(C)** The number of HLA alleles varies greatly with each typing resolution. In this figure, “pseudo-sequence” refers to the amino acid residues directly involved with the binding of the peptide as shown in **D)**. Each of the typing resolutions shown in this figure are subgroups of the higher typing resolutions, meaning that it is always possible to convert unambiguously from e.g. 2-field resolution to P group resolution. Note, that null alleles are disregarded at P group and pseudo-sequence resolution, as these do not correspond to an expressed peptide sequence. The data is acquired from the IPD-IMGT/HLA database (1) release 3.48.0. **(D)** A binding pocket of an HLA class II molecule (HLA-DR). The residues directly involved in peptide binding (the pseudo-sequence) are highlighted in purple. DRA is shown in green and DRB1 in blue and a melanoma antigen in the binding pocket is shown in red. Protein data was obtained from the Protein Data Bank (10, 11) and the figure was made using PyMOL (**?**) Figures **(A)** to **(D)** were crated using https://www.diagrams.net/.

Cells can present peptides to T-cells both from pathogens (non-self peptides), tumor mutations (neopeptides) and cells native to the body (self-peptides). T-cells are however generally able to recognise the difference. This means that a cell displaying a peptide, thereby indicating that it is infected by a virus or developing into a tumor cell, can trigger an immune response that a healthy cell would otherwise avoid (3, 4). The binding of a peptide to HLA and subsequently to a T-cell receptor (TCR) is highly specific, and a given HLA molecule will only bind and display a peptide if it matches the HLA molecule’s binding cleft, which is also known as the antigen recognition domain (ARD) (5, 6). The HLA region is one of the most polymorphic regions in the human genome with over 33000 known allele sequences as of writing (IPD-IMGT/HLA Release 3.48.0). The most polymorphic HLA genes are the classical class I genes, HLA-A, -B and -C as well as the class II genes HLA-DRB1, -DQB1 and -DPB1 (1). Due to the large number of alleles, the naming of specific HLA alleles follows a special naming convention as illustrated in figure 1**(B)** and **(C)** (7, 8). HLA nomenclature is comprised of four fields, each made up of 2-3 digits and each describes a specific allele with increasing precision.

The most important part of the HLA molecule is ARD, which is encoded by exons 2 and 3 in class I molecules and by exon 2 in HLA class II molecules. The most important differences between alleles are therefore the ones affecting the nucleotides in this region (7). Two official ARD based HLA typing resolutions exist: G group resolution which clusters alleles with identical nucleotide sequences in ARD coding exons and P group resolution which groups alleles with identical peptide sequences in the ARD. An overview of these can be found at^1^. A 2019 article (12) argued that mismatches outside the ARD are for the most part not important and focusing on them could delay or prevent organ transplantation. The recommendation was therefore that “clinical decision-making should focus only on the sequence of the antigen recognition domain (ARD) with the exception of common non-expressed alleles that are distinguished by variation outside of the ARD”. This recommendation is followed by using P group resolution and accounting for null alleles separately. Alleles can be further grouped based on the residues that are directly involved with the binding of the peptide to the HLA molecule (see figure 1 **D**). This grouping method is used in tools predicting peptide-HLA binding such as NetMHC-pan4.1, but is not an official typing resolution (13).

HLA typing is the process of determining an individual’s specific HLA alleles. HLA typing is used widely since the peptide presentation is a crucial part of the adaptive immune system and depends on the specific HLA allele. Some examples include the study and prognosis of infectious diseases, autoimmune diseases and cancer, as well as the discovery of neoantigens in cancer treatment and for finding compatible donors for organ transplants (5, 14, 15).

Allele typing in the HLA region is significantly more difficult than allele typing in other regions. This is mainly due to the high degree of polymorphisms but also the high degree of sequence homology between both different alleles and between alleles of different HLA genes. HLA genes are further co-dominantly expressed, giving an enormous amount of possible HLA profiles (16).

Traditional HLA typing uses lab-based methods which can be slow and expensive and often generates data only for the purpose of HLA typing. The rapid development of next-generation-sequencing (NGS) has however resulted in large amounts of easily available sequencing data which can be used for HLA typing by using recently developed computational tools (17).

NGS based HLA typing tools can roughly be divided into two groups - those using *de novo* assembly-based methods and those which directly align to a reference sequence. The alignment-based methods either use a traditional linear reference or a graph-based reference / graph-based alignment algorithm (18, 19). The tools further differ on which HLA genes they can type and on the sequencing data, which they use for typing.

A 2019 review of NGS-based HLA typing noted the lack of systematic benchmarking of the many available HLA typing algorithms (18) and although several benchmarking studies have been published there does not seem to be one best performing tool, instead the performance depends on the sequencing data and the HLA gene (20, 21). Some tools, such as Optitype (17) and Polysolver (22) only offer typing of class I genes, and the developers of the graph-based method Kourami write, that the tool is developed for high-coverage WGS data (23).

New alleles are registered and named by a World Health Organisation committee and stored in the IPD-IMGT/HLA database. The database is continuously updated and errors are corrected but the database is not complete. New alleles are still being discovered and the full genomic sequence is not known for all registered alleles. Some entries are still missing the non-ARD coding exons and/or the introns (1). The application of NGS-based HLA typing is not limited to presently living individuals but has also been used in studies of ancient genomes for example to find specific HLA alleles that increase susceptibility or protection to a specific disease. Sequencing of ancient DNA (aDNA) is often limited by a low depth of coverage, short DNA sequence length and chemical damage to the DNA (24). However, studies of aDNA have still used modern HLA typing tools such as Optitype (25) and an adaptation of HLAssign (26) to perform HLA typing on ancient individuals. HLAssign relies on data generated using targeted HLA enrichment (27), while Optitype is designed for general, non-enriched sequencing data (17). (28) noted, that although these methods have been used for HLA typing from aDNA, there is no structured study of their expected performance on this type of data.

In this study, we present a comprehensive review of the performance of freely available HLA typing methods based on NGS. Specifically, the five HLA genes HLA-A, -B, -C, -DRB1 and -DQB1 are typed using whole-exome sequencing (WES) data, as this type of data and these loci are widely used in clinical settings (29, 30). With this analysis, we demonstrate the first use of P group resolution and pseudo-sequence resolution in a benchmarking study of WES based HLA typing.

We find that HLA*LA had the highest typing accuracy across the five HLA loci, and that the typing resolution does not have an effect on which tool performed the best. We show that the impact of the sequencing coverage on the HLA typing accuracy depends heavily on both the tool and the HLA locus. A depth of coverage of at least 100X is advisable for accurate typing of all five HLA genes - even for the best performing tools.

Additionally, we estimate Optitype’s performance on aDNA by running it on a simulated dataset that mimics aDNA samples in terms of coverage, read length and adding simulated chemical damage. Interestingly, we find that read length does not matter as much as the depth of coverage however, Optitype requires a coverage between 10X and 20X to achieve a typing accuracy of above 90 % which is often prohibitively high for ancient DNA samples.

## Materials and Methods

### Selection of HLA typing tools

There are numerous NGS based HLA typing tools available. This study focused on freely available tools which ran on WES data and have shown promising results in previous benchmarking or proof of concept studies. This means that tools such as HLA-HD, Polysolver and OncoHLA, which require some form of license, were not included in this study (22, 31, 32). The final selection of tools is listed in table 1. STC-Seq was included as a reference to illustrate how a more simple algorithm, which is designed for HLA enriched data, perform on lower coverage WES data. Optitype was downloaded from^2^. CBC 2.9.5 was used as ILP solver as it was found to be more stable than CPLEX 12.7 which often did not converge to a solution. Kourami was downloaded from^3^, HLA*LA was downloaded from^4^, HISAT-genotype using its web-guide at^5^ and STC-Seq was downloaded from the BioCode website^6^.

**Table 1.**
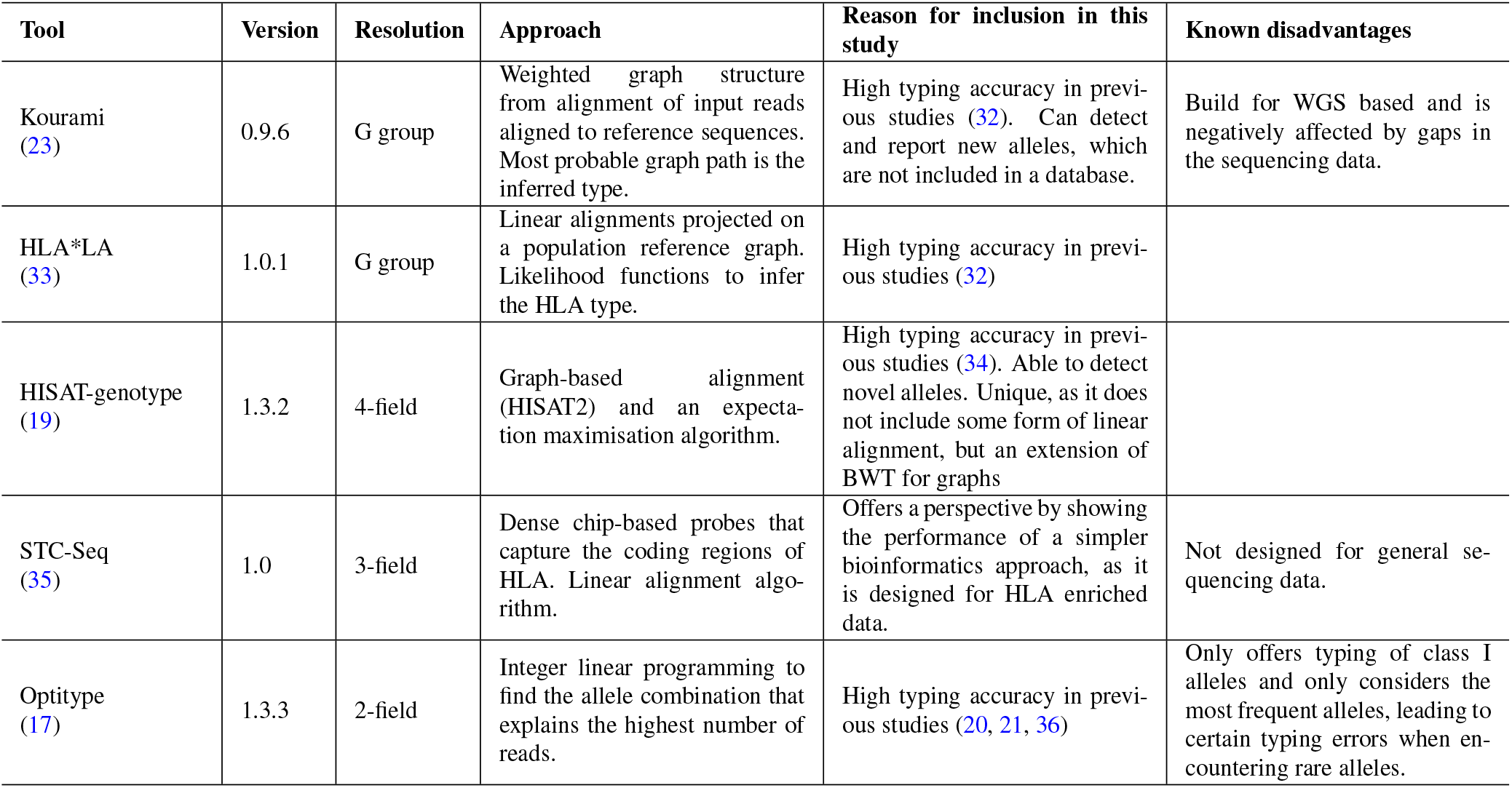
Details of the five HLA typing algorithms included in this project. Each of the original articles describing the tools contains some form of benchmarking study that demonstrates the capabilities of the tool.

All tools in the analysis were given 10 threads and as much memory as they needed.

### Benchmarking dataset

To evaluate the performance of the HLA typing tools, we used a reference dataset consisting of WES samples from 829 individuals taken from the 1000 genomes phase 3 dataset (37). The sequencing coverage of the samples ranged from 37X to 456X with the median being 86X. The true HLA types of the samples was determined using laboratory based methods. 819 of the 829 samples were typed in a 2014 study (38) and further validated in a 2018 study (39). Some alleles were in 2018 found to have been mistyped in 2014, and these were updated in our dataset. The last 10 of the 829 samples were typed by (40) and further validated by (39). We found that this dataset contained some alleles, whose names have changed since 2014. For both the 1000G dataset and for the predictions made by the HLA typing tools, these alleles were converted to their newest name. A full overview of all deleted/renamed alleles can be found at ^7^. This constructed dataset is referred to as the 1000G dataset in the remainder of the paper.

For the majority of the individuals in the 1000G dataset, the HLA typing was only available in 2-field resolution and the tools could therefore only be evaluated at 2-field resolution or lower. For some individuals, the HLA typing in the dataset is ambiguous. One example is the individual NA12287 with HLA-B typing: 15:01/15:03 and 15:01/15:26/15:12/15:19. In these cases, predictions by the tools were counted as correct if they matched any combination of correct alleles. Some of the samples in this dataset were also used in the development of the HLA typing tools or at least included in the proof of concept study in the original articles introducing the tools. Out of the 829 individuals, 31 were included in the paper introducing Kourami, 28 for HLA*LA and 95 in the paper introducing Optitype.

### Performance evaluation and typing resolution

The five HLA typing tools were evaluated on several different metrics with the most important being the typing accuracy, which is the number of correct predictions out of the total amount of HLA alleles. The typing accuracy was found for the resolutions: 1-field, pseudo-sequence, P group and 2-field (see figure 1). Some tools did not return a prediction for all alleles, and the individual tools’ call rate (number of predictions a tool returned as a fraction of total number of alleles) was therefore also noted. The time (CPU and real), as well as the memory use of each HLA typing tool, was registered and the full pipeline for running the tools is illustrated in figure S1 in the supplementary material.

### Typing resolution

The reference HLA alleles in the 1000G dataset are in 2-field resolution which cannot be unambiguously converted to G group resolution. This is because 2-field resolution separates alleles based on differences in the full amino acid sequence while G group resolution separate alleles based on genomic differences in the ARD coding exons. In this study, predictions in G group resolution were converted to “2-field” resolution by trimming the third field, but this is not a perfect approach, as is discussed in section.

The pseudosequence resolution, which for HLA-DRB and - DRA is shown in figure 1, was presented by (41) and we use the same approach to constructing the pseudo-sequences as was described in the original article. That is, a pseudo-sequence consists of the 34 amino acids, which are within 4 Å of a peptide bound to the HLA molecule. Alleles which share these 34 amino acids belong to the same pseudo-sequence group.

### Downsampling

The 1000G dataset described in section has 230 samples with a coverage of at least 100X. These samples were included in a downsampling study to investigate how the performance of each tool depended on the coverage of the input sample. The performance of the tools was examined for each of the the 230 samples at depths of coverage of 1X, 2X, 5X, 10X, 20X, 50X, 75X and 100X. Downsampling was performed by first finding the full sequencing depth of the CRAM files used in the 1000G dataset. This was done using *mosdepth* (version 0.2.6) (42). Hereafter, alignment files containing potential HLA reads were downsampled using *samtools view -s* to achieve the range of needed coverages (43). The resulting files were then HLA-typed in the same way, as in the main study (see supplementary material figure S1)

### Optitype’s performance on simulated ancient DNA

The 50 WES samples with the highest depth of coverage from the 1000G dataset were used to simulate an aDNA dataset and evaluate Optitype’s performance on this type of data. The downsampling procedure to specific sequencing depths was the same as the one described in the downsampling analysis. Gargammel (v. 1.1.2) (44) was used for trimming reads to specific read lengths and for the addition of simulated chemical damage. The added chemical damage was done in a way so as to simulate the chemical damage in the samples from a 2021 study of medieval plague victims(45). Specifically, the added damage was 20 base of C to T substitutions after the 5’ end, the substitution rate for the base immediately after the 5’ end was 7% and fell below 1% 3 bases after the 5’ end. There were also 20 bases before the 3’ end of G to A substitutions, the base right before the 3’ end has a substitution rate of 6% and then fell below 1% 3 bases before the 3’ end. Reads were downsampled to the coverages 1X, 2X, 5X, 10X, 20X and 50X and read lengths were varied over 10, 13, 15, 20, 25, 30, 35, 45, 55 and 65 base pairs. The WES samples did not have enough reads to achieve a coverage of 50X for the lower read lengths, so some read length/coverage combinations could not be performed.

## Results

The typing accuracy for the HLA typing tools was measured in four different typing resolutions, 1-field, pseudo-sequence, P group and 2-field and the results presented in this section is available in all four typing resolutions as supplementary data. The focus will however primarily be on the clinically relevant P group resolution.

### Overall performance of the tools

Figure 2 outlines the performance of the five HLA typing tools on the full 1000G dataset across four different typing resolutions. HLA*LA, Optitype and HISAT-genotype all have a call rate of 100% across the genes, that they offer predictions for. Kourami fails to return a prediction for almost 9% of alleles and STC-Seq for more than 30%. In Kourami’s case, a failure to return a prediction often happens when there is not enough data to sufficiently cover important regions leading to Kourami’s graph structure being disconnected (23). Optitype is the best performing of the tools for class I genes and has a typing accuracy close to 100% across all typing resolutions. These results match the results of previous studies evaluating the performance of Optitype (20, 21). HLA*LA is the best performing tool for the two HLA class II genes as well as across all five HLA genes.

**Fig. 2.**
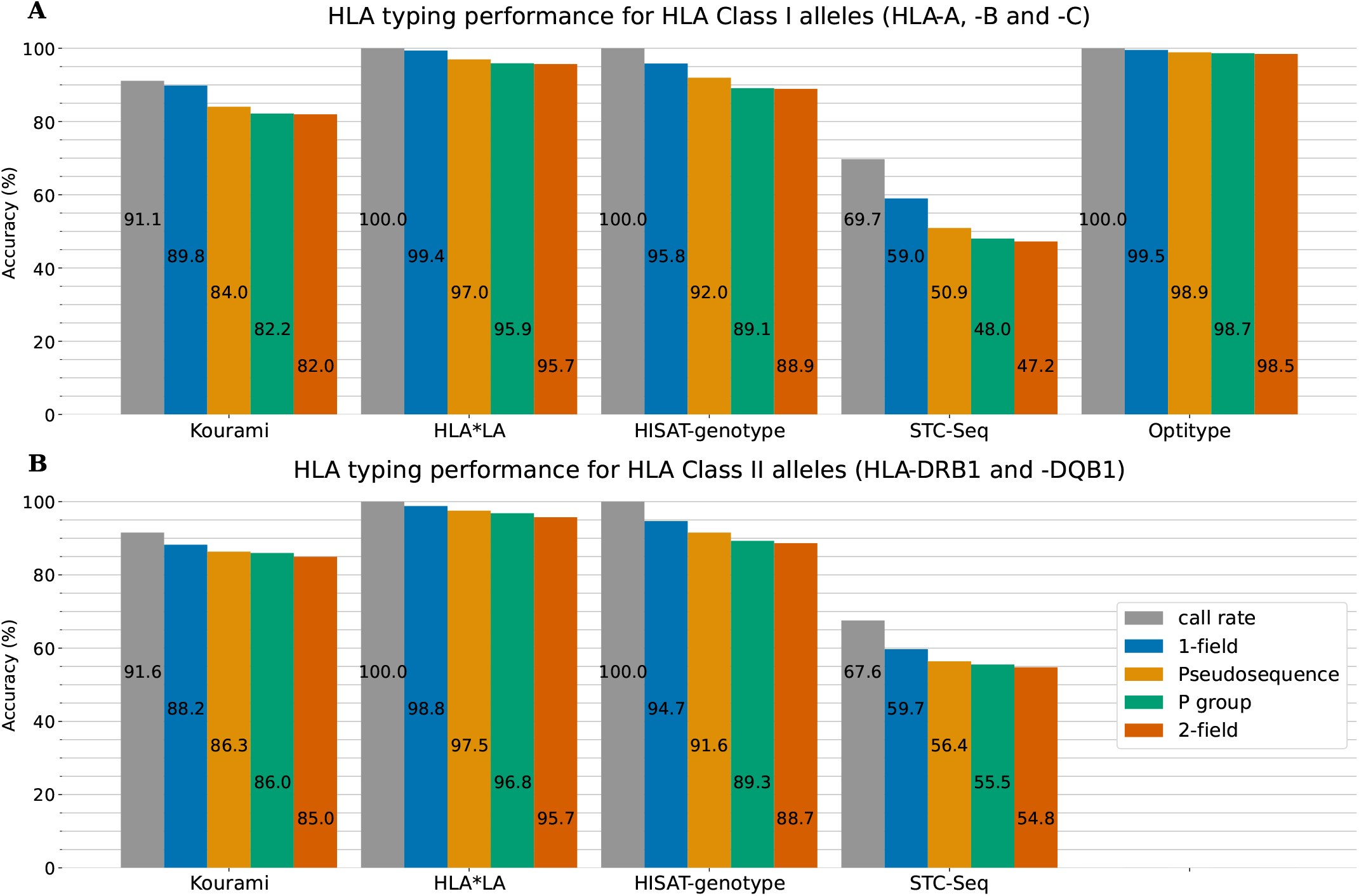
The five HLA typing tools’ typing accuracy (number of correctly called alleles out of the total amount of alleles) and call rate (number of called alleles out of all alleles) in 1-field, pseudo-sequence, P group and 2-field resolution for the three HLA class I genes HLA-A, -B and -C **(A)** and the two HLA class II genes HLA-DRB1 and -DQB1 **(B)**. Optitype does not offer class II typing and is therefore not listed in (B). The full results, stratified on the individual HLA genes, can be found in supplementary figure S2

HLA*LA performs almost as well as Optitype at 1-field resolution, while the difference in performance between the two tools is larger at higher typing resolutions. In 2-field resolution, HLA*LA on average mistypes one out of 24 class I alleles (one mistyping per 4 individuals with 6 alleles each), which is thrice as many as Optitype.

For both HLA class I and class II genes, the typing resolution does not change which tools perform the best. Across all typing resolutions and class I genes, Optitype has the highest typing accuracy, HLA*LA has the second-highest followed by HISAT-genotype, Kourami and STC-Seq in that order. The order is, save for Optitype, the same across the class II genes. Using P group resolution instead of 2-field does however make some difference to the typing accuracy. HLA*LA miscalls 141 out of 3316 class II alleles in 2-field resolution, but 36 of the 141 are correct calls in P group resolution and 59 of the 141 are correct calls in pseudo-sequence resolution. Figure 3 shows the distribution of the peak memory usage and real-time usage across the 829 samples in the 1000G dataset for each of the five tools included in this study. Generally, STC-seq and Optitype and Kourami use the least memory per sample, with median usages of 0.38 GB, 1.1 GB and 1.7 GB respectively. For a few of the samples, STC-Seq and Optitype use more than 8GB of memory, while the most memory demanding sample takes 6.4 GB for Kourami. HISAT-genotype uses 8GB of memory for almost all samples, indicating that this is a built-in restriction. Allowing HISAT-genotype to use more than 8GB of memory could perhaps reduce the runtime of the tool. HLA*LA uses by far the most memory with a median of 31 GB per sample and the most memory demanding sample (NA18504) requiring over 600 GB of memory. The high memory usage is due to HLA*LA’s expensive alignment step that uses dynamic programming (23) and is also noted in an HLA*LA blogpost^8^.

**Fig. 3.**
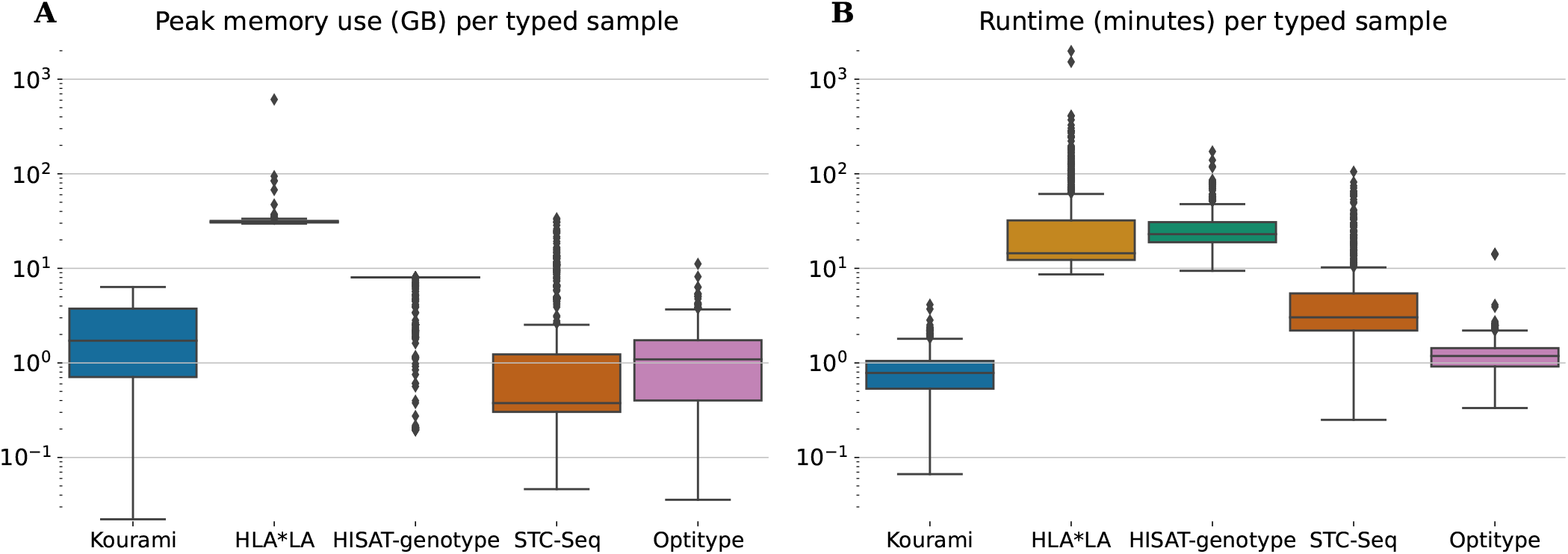
**(A)** The peak memory usage and **(B)** real-time usage of HLA typing for each sample in the full 1000G dataset. Note, that this is only for the tool-specific HLA typing step and therefore does not include the previous steps such as the extraction of HLA reads.

HLA*LA and HISAT-genotype spend the most time per typed sample. HISAT-genotype’s median time (23 minutes) is higher than HLA*LA’s median time usage (15 minutes) but HLA*LA spends more than a day for a few samples, whereas HISAT-genotype at most spends 172 minutes. STC-seq spends more than an hour for some samples, but types most samples in under 10 minutes. Kourami and Optitype type most samples in less than 2 minutes. The CPU time usage of the tools can be found in figure S3.

Kourami, Optitype, HISAT-genotype and STC-seq can, all be run on a system with less than 16GB of memory and run most samples in less than an hour with Optitype and Kourami generally requiring far less time. HLA*LA requires much more memory than the other tools and spends more than an hour for 129 samples and more than 24 hours for 2 samples. These high resource requirements should be kept in mind when choosing this tool.

### Downsampling analysis

The typing accuracies presented in this section are, unless specified, all in P group resolution and therefore match the results shown in figure 4. This figure shows that the typing accuracies of Kourami, HLA*LA, Optitype, HISAT-genotype and STC-Seq depend highly on the coverage of the samples when the coverage varies between 1X and 100X. A higher coverage correlates with a higher typing accuracy but this correlation is not linear and differs between the HLA typing tools. Some HLA typing tools maintain a high typing accuracy when the depth of coverage decreases, while others require a high coverage for accurate typing. Optitype performs the best for low coverage samples and its typing accuracy only drops below 90% when coverage is below 20X. HLA*LA, which performs almost as well as Optitype at 100X, has a typing accuracy of 72.7% at 20X for the class I genes. STC-Seq and Kourami are both reported to have a typing accuracy very close to 100% when typing from each tool’s preferred sequencing data (HLA enriched data for STC-Seq and high coverage WGS data for Kourami) (23, 35), but as is shown in this study, the tools do not perform well on low coverage WES data.

**Fig. 4.**
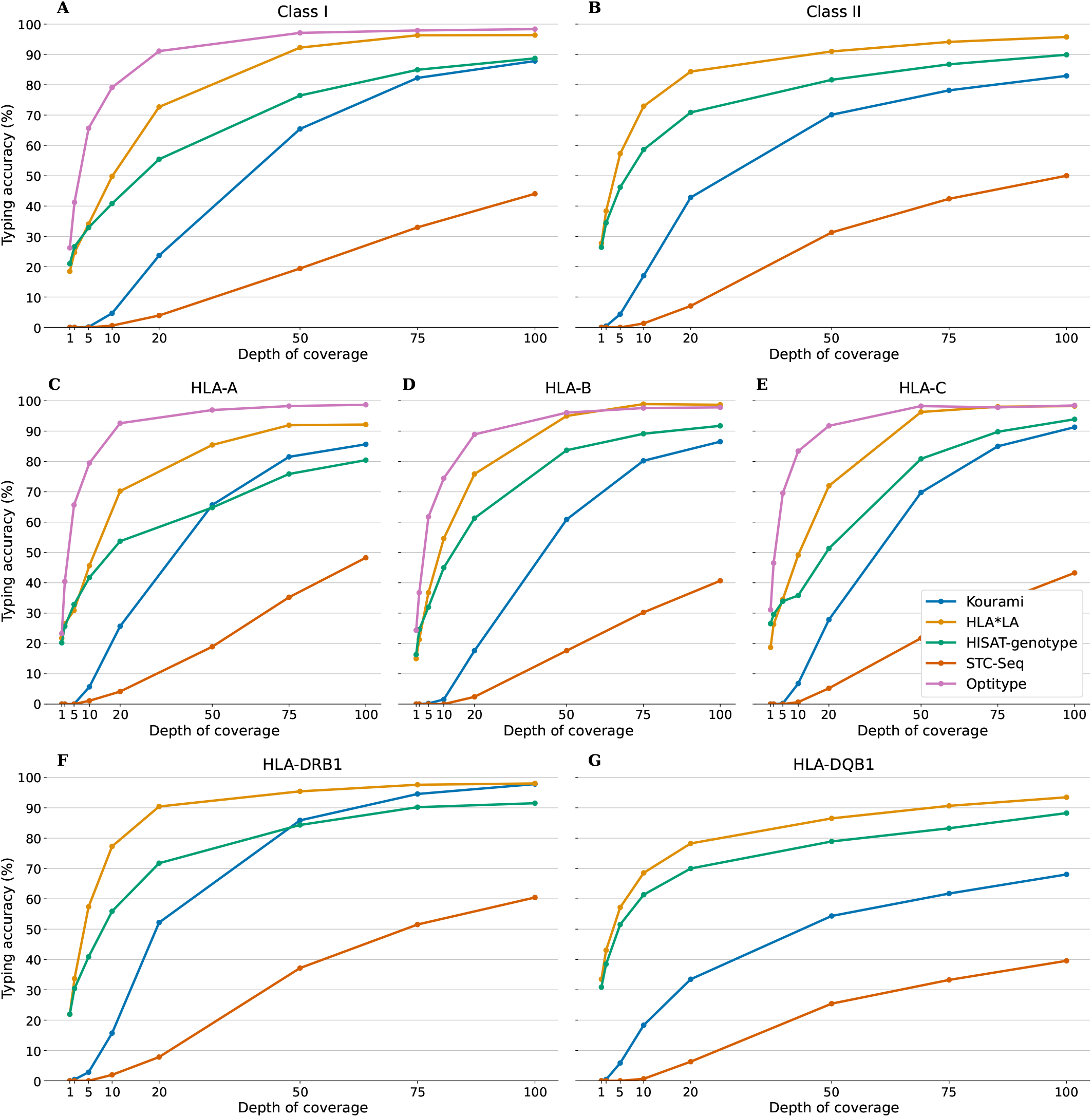
The typing accuracy of Kourami, HLA*LA, Optitype, HISAT-genotype and STC-Seq in P group resolution for 230 WES samples across coverages ranging from 1X to 100X. The top two panels (**(A)** and **(B)** show the overall typing accuracy for HLA class I and II, while figures **(C)** to **(F)** show the performance for each individual HLA gene. The performances of the tools are also available in 1-field, 2-field and pseudo-sequence resolution in as supplementary data

Figure 4 **(C)** to **(F)** show that the HLA typing accuracy not only varies between tools and with the depth of coverage but also between HLA genes. HLA typing tools in this study achieve a higher typing accuracy for HLA-DRB1 than they do for HLA-DQB1. At 100X HLA*LA has a higher typing accuracy than Optitype for HLA-B and HISAT-genotype is much better at typing HLA-B and -C than HLA-A. The difference in performance between HLA-DRB1 and HLA-DQB1 is especially prevalent for Kourami. At 100X coverage Kourami has a call rate of 100% and a typing accuracy of 97.8% for for HLA-DRB1 while for DQB1 the tool only has a call rate of 78.2% and a typing accuracy of 68.0%.

The gain in typing accuracy from increasing the depth of coverage is, as expected, generally larger when the coverage is low, and there are diminishing returns from an increase in coverage, when the coverage is already high. This trend, however, depends on the HLA typing tool and the HLA gene. For the HLA class I genes, Optitype and HLA*LA do not benefit much from an increase in coverage from 75X to 100X, while the remaining three tools might even benefit from increasing the coverage beyond 100X. For HLA-DRB1, HLA*LA and HISAT-genotype’s perform almost equally well at 75X and at 100X, but for HLA-DQB1, the tools perform notably better at 100X than at 75X. HISAT-genotypes typing accuracy even increases more from 75X to 100X than it does from 50X to 75X.

The findings from the downsampling dataset generally agree with those from the full dataset, but there are some notable differences. For the full dataset, Optitype has the highest typing accuracy for all three class I genes (see figure S2), but for the downsampling, HLA*LA outperforms Optitype for HLA-B and is as good as Optitype for HLA-C for coverages of 75X and above. Kourami has a higher typing accuracy than HISAT-genotype for HLA-A on the full dataset, but the downsampling shows that HISAT-genotype has a higher typing accuracy when the coverage is below 50X. Kourami also shows a large improvement with an increase in coverage for HLA-DRB1 and at 100X it performs almost as well as HLA*LA.

### The impact of short fragments and DNA damage

Figure 5 shows Optitype’s expected performance across varying coverages, read lengths and with damage added to the reads. The figure only shows a subset of the results from the combinations between coverage/read length and added DNA damage, but the full results are available as supplementary material. We found that Optitype was unable to return any results when the read length was 10, regardless of the coverage. As also shown in figure 4, Optitype’s typing accuracy depends largely on the coverage, when this varies between 1X and 20X. At lower coverages, the read length seems to have some influence on typing accuracy, but this effect vanishes when the coverage increases. Between 1X and 10X, Optitype performs the best when the read length is at 45 and both lower and higher read length result in a drop in typing accuracy. There is still a slight performance gain by increasing the coverage from 20X to 50X, as shown in supplementary figure S5, but at this stage, an increase in read length does (as long as it is above a minimum of around 25) not result in a higher typing accuracy.

**Fig. 5.**
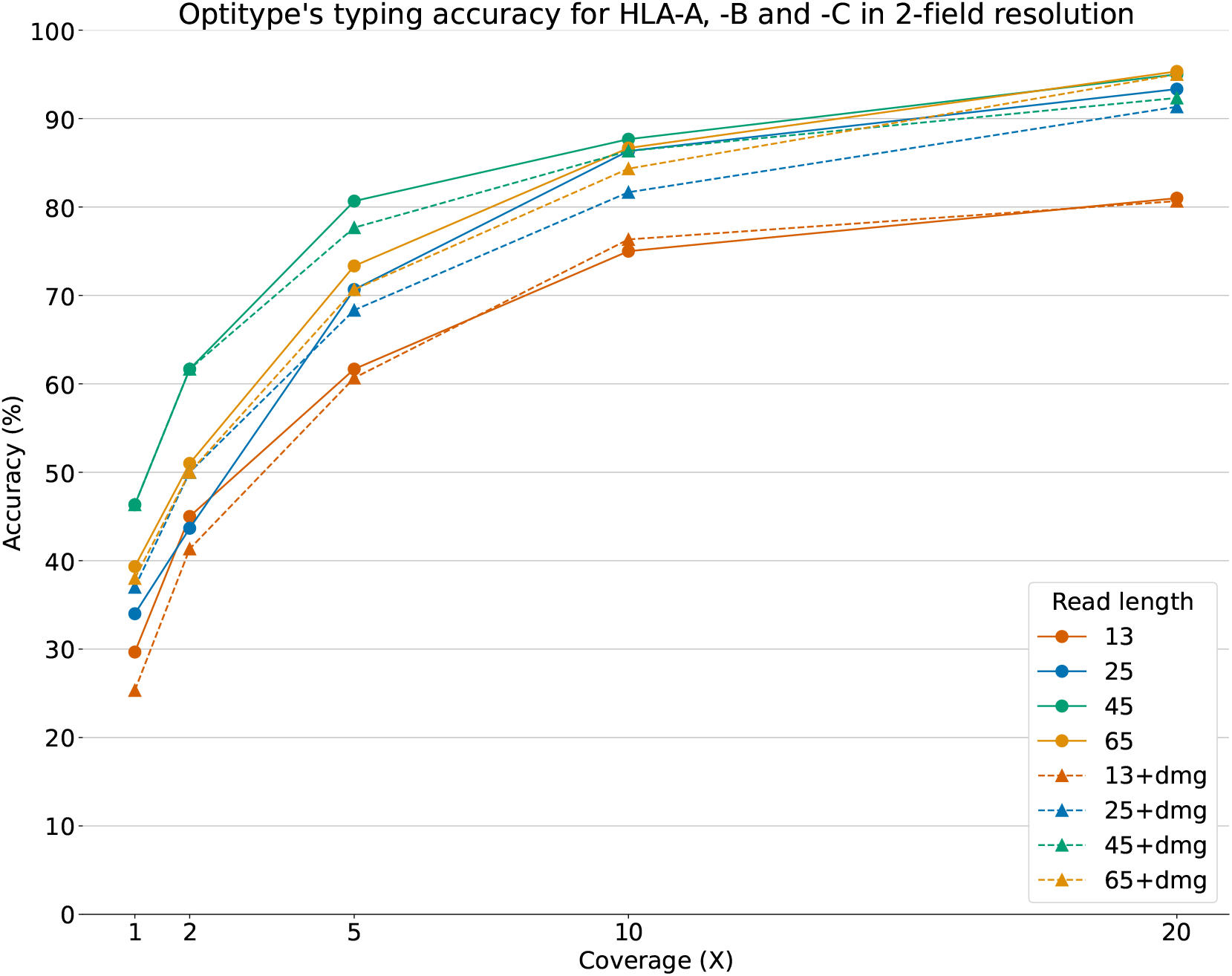
Optitype’s typing accuracy for HLA-A, -B and -C across 50 samples and across varying read lengths, coverages and with/without artificially added chemical damage. The typing accuracy depends primarily on the depth of coverage. Note, that the downsampling, read cutting and addition of DNA damage was done after an extraction of HLA reads. This figure therefore represents a best case scenario, where assigning reads to the HLA region is as accurate for short reads as it is for the full length reads.

The addition of simulated DNA damage did not notably impact the typing accuracy at any coverage or read length. The typing accuracy in figure 5 and supplementary figure S5 is shown in 2-field resolution, as this resolution was used in (25).

## Discussion

This study investigated the performance of five NGS based HLA typing tools: Kourami, HLA*LA, HISAT-genotype, Optitype and STC-Seq for the five HLA genes: HLA-A, -B, -C, -DRB1 and DQB1. The tools were evaluated on 829 WES samples from the 1000 genomes dataset as well as on a downsampled, subset of these to evaluate the impact of coverage on typing accuracy. The typing accuracy was evaluated at four different typing resolutions, 1-field, 2-field, P group and pseudo-sequence, with the two latter not having been explored in previous studies of the performance of HLA typing algorithms.

HLA*LA was found to have the highest overall typing accuracy (96.3% in P group resolution for the full dataset) and the highest typing resolution for the two HLA class II genes (96.8% in P group resolution) while Optitype was found to have the highest typing accuracy for the three HLA class I genes (98.7% in P group resolution). The tools varied greatly in computational resource consumption with HLA*LA requiring 30 GB of memory and often more than an hour per typed sample whereas Optitype only required 1 GB of memory and rarely more than a couple of minutes.

Evaluating the HLA typing on samples across varying coverages showed, that the typing accuracy of all the tools depended greatly on the depth of coverage, although this dependency differed between both the HLA typing tools and across the five HLA genes. A sequencing depth of 50X was enough for class I typing using Optitype, while accurate typing of HLA-DQB1 required at least a sequencing depth 100X even for the best performing tool, HLA*LA.

### Ambiguous typing results

In this study, we converted predictions from HLA*LA and Kourami in G group resolution to 2-field resolution by simply removing the third field. However, converting predictions both from G group to 2-field and vice versa is ambiguous, as G group resolution focuses on the genomic sequence of the ARD region, while 2-field resolution focuses on the peptide sequence of the full protein.

There are examples of proteins (alleles in 2-field resolution), which are part of two G groups that differ at the second field. An example is HLA-C*02:02, as this could both be C*02:02:02:01 (which belongs to the C*02:02:02G), C*02:02:01 (which is not part of a G group) or even C*02:02:37 (which is part of the C02:10:01G group that differs from C*02:02 at the second field).

Conversely, there are examples of G groups containing alleles, that differ in 2-field resolution. The G group HLA-A*01:01:01G contains A*01:01, but also A*01:32 and 78 other alleles that differ in 2-field resolution. This ambiguity poses a problem when using typing methods such as HLA*LA and Kourami, which return the results in G group resolution, but also when evaluating their performance on the 1000G dataset in 2-field resolution. If the correct allele was HLA-A*01:32, the correct prediction in G group resolution would be A*01:01:01G, but should this prediction be counted as correct in 2-field resolution even though the prediction could refer to more than 70 individual alleles in 2-field resolution? One approach to this ambiguity is to convert both the reference alleles and all predictions to G group (e.g. HLA-A*01:01:01G) and then trim to 2-field resolution (HLA-A*01:01), but this can not be done unambiguously for alleles such as C*02:02, as described previously. A similar approach, that partly solves the ambiguity issue is to convert predictions to P group resolution. 2-field and G group resolution can, save for null alleles, both unambiguously be converted to P group resolution.

The approach used in this study to convert G group predictions to 2-field resolution allowed for a somewhat fair tool comparison in 2-field resolution, but it is not as good a comparison as in P group resolution.

A whole other factor, which impacts the accuracy of the results in 2-field resolution is that the experimental method for discovering the true HLA types of the 1000G dataset often only focus on the ARD coding regions and does not sequence non ARD coding exons (38). This ambiguity is found in the 1000G truth set. There are some cases, where e.g. a true allele is noted solely as C*17:01, but the true allele might be any allele in the C*17:01P group. An HLA typing tool calling the allele as C*17:02 (which is part of the C*17:01P group) will then unfairly be noted as having mistyped the allele. Like the problem with G group/2-field conversion, this problem can also be solved by converting both the 1000G dataset and the predictions to P group resolution.

### The impact of depth of coverage

A 2013 study (46) stated that the depth of coverage of many of the samples in the 1000 genomes dataset, was too low for HLA typing. Another article from 2018 disputed this statement and stated that they found no correlation between typing accuracy and depth of coverage although noting that a minimal coverage was required (20). The results outlined in figures 2 and 4 clearly show, that even the samples with the lowest coverage in the dataset (between 35X and 40X) can be typed accurately, if a tool suited for low coverage samples is used, effectively disproving the statement from (46). The results also show that the typing accuracy depends strongly on the coverage of the sample, in contrast to what is stated in the 2018 article, and further that a “required minimal coverage for optimal typing” depends on both the HLA typing tool and the HLA gene.

Optitype and HLA*LA’s overall performances do not improve much when the coverage is increased from 75X to 100X, but the other three tools do and they might even see an improvement if the coverage is increased beyond 100X. Achieving a typing accuracy above 90% for HLA-DQB1 also requires WES coverage of at least 100X, even for the best performing tool, HLA*LA. These results align roughly with minimal coverage recommendations for clinical WES of 120X (47) and WES based HLA typing of 100X (34). The results from this study are however more detailed and clearly indicate that for some genes e.g. HLA-DQB1 or tools e.g. Kourami, it is even beneficial with coverage above 100X. The coverage of the samples in the 1000G dataset varies between 37X and 456X, with many samples having a lower coverage, than what is required for optimal typing. The tools’ performance on this dataset might therefore not be an accurate estimate of the performances on clinical WES with coverage above 100X. A better estimate of this could be the downsampling results, specifically the performance of the samples at 100X. This is a redeeming factor for e.g. Kourami, which had a mediocre performance on the full dataset, but where the results of the downsampling study show, that the lower coverage of the full dataset likely impaired Kourami’s performance much more than it did Optitype’s.

HLA*LA is the best performing tool on average, across the five HLA alleles but the tool does require an extensive amount of memory and is relatively slow. Mistypings/mismatches at HLA-DQB1 is preferred compared to other HLA genes (48), and a lighter and faster alternative to HLA*LA is, therefore, Optitype for class I typing and, assuming coverage of at least 100X, Kourami for class II typing. For the 230 high coverage samples at 100X, Optitype had a P group typing accuracy of 98.3% across the three class I genes (HLA*LA had 96.4%) and Kourami had a P group typing accuracy of 97.8% for HLA-DRB1 (HLA*LA had 98.0%).

### Allele diversity of HLA-DQB1

All four tools, which offers class II typing, perform worse for HLA-DQB1 than for HLA-DRB1. This could indicate, that HLA-DQB1 typing is more difficult, but there could also be other explanations of this. (49) found that HLA calls of the 1000 genomes dataset were biased towards reference alleles, meaning that the 1000G dataset contains an unexpectedly high allele frequency of HLA alleles similar to the reference genome used in the 1000 genomes project and conversely an unexpectedly low frequency of alleles which differed to the reference. This effect was found for HLA-A, -B, and DQB1, to some degree for HLA-C but not for HLA-DRB1. Another explanation to the HLA typing tools’ performance difference between DRB1 and DQB1 could therefore be, that the 1000G dataset contains more miscalls for DQB1 than for DRB1 and that some of the “miscalls” by the HLA typing tools are actually correct.

Figure S7 shows the number of unique allele calls made by each tool as well as the number of unique alleles in the 1000G dataset across the five HLA loci. The 230 individuals included in the downsampling study only share 16 unique HLA-DQB1 alleles according to the 1000G dataset while the three best performing HLA typing tools, that provide typing for HLA-DQB1 (Kourami, HISAT-genotype and HLA*LA), on average call more than twice as many. For DRB1, this difference is much smaller. The 230 individuals share 42 unique alleles according to the 1000G dataset, while the three HLA typing tools on average call 51 unique alleles. The comparatively few unique calls for HLA-DQB1 could support the previous indication, that the HLA-DQB1 sequencing was biased towards the reference sequences, although it could also be due to other factors, such as the higher overall diversity of HLA-DRB1 compared to HLA-DQB1 (1)

Figure 4 **(G)** shows one trend, which could indicate, that the HLA-DQB1 typing in the 1000G dataset is correct. The HLA typing tools, which provide HLA-DQB1 typing, do not seem to hit a performance ceiling, but continue to improve, when the coverage is increased. If a large amount of the reference HLA-DQB1 alleles were wrong, an increase in coverage would likely not lead to an increase in typing accuracy.

### Optitype’s performance on ancient DNA

With figure 5, we present the first benchmarking of HLA typing tools on aDNA. The figure shows, that the coverage is the most determining factor for a high typing accuracy, but that at low coverages, both a too high and a too low read length can impair typing accuracy. It is expected, that a low read length negatively affects allele typing - especially for a highly polymorphic region, such as the HLA region. However, it is surprising that the typing accuracy for the samples with a read length of 65 is lower than when the read length is 45. One reason for this might be, that Optitype’s typing algorithm consists of creating a binary hit matrix where the predicted HLA alleles are the ones explaining the highest *number* of reads. When the read length is increased, but coverage is kept constant, the number of reads decreases. The sample with reads of length 45, therefore, contains more reads than the one with a read length of 65 at the same coverage as is illustrated in supplementary figure S6. The difference in performance, which can be attributed to read length, decreases at higher coverages and the typing accuracy is almost the same at a depth of coverage of 10X. The addition of DNA damage did not impair the performance notably. However, the DNA damage applied in this study corresponded to that on specific samples from the 16th century and e.g. older samples or samples from individuals stored in different conditions can have a larger degree of DNA damage, which could have a bigger effect on the accuracy of the HLA typing.

Studies of aDNA often use methods designed for contemporary data for variant calling, which can lead to inaccurate results (50). (25) performed a genomic analysis of individuals who lived around 3200BCE and part of this was an HLA analysis of 23 individuals where they observed some “striking shifts in allele frequencies”. The study used Optitype for HLA typing in combination with another method but did so without investigating Optitype’s expected performance at the coverages found in their aDNA dataset. The median coverage of the samples, which were HLA typed in the study ranged from 0.07X to 18.2X with the median being 4.3X. The average read length spanned from 51.8 to 67.6 with the median being 58.2. Our results outlined in figure 5 show, that most of the samples included in (25) had such low coverage, that Optitype likely only returned a correct prediction for little over half of the alleles. The study did not rely solely upon Optitype for HLA typing, but our findings show, that having a high coverage is crucial and that Optitype’s typing results are not reliable for low coverage aDNA samples.

## Author Contributions

NT, MK, SG, TT and GR conceived this study. NT gathered the data, installed and ran the tools, performed the subsequent data analysis and wrote the manuscript. MK and GR provided support to the data analysis, GR assisted with the analyses related to ancient DNA. All authors reviewed and provided feedback on the manuscript.

This study is based on the NT’s Master’s thesis (51), which can be found at https://findit.dtu.dk/en/catalog/60339f5ad9001d01650f4d5d

## Acknowledgments

The authors want to thank the team behind Denmark’s National Life Science Supercomputing Center (Computerome - https://www.computerome.dk/) for providing the computational resources for this project and Christian Garde from Evaxion Biotech for a variety of help throughout the project.

## Data Availability Statement

The 1000 genomes WES data as well as the HLA typing is available at: https://www.internationalgenome.org/ The HLA typing of the 10 samples, which were typed as part of the study introducing ATHLATES (40), can be found at: https://www.ncbi.nlm.nih.gov/pmc/articles/PMC3737559/bin/supp_gkt481_nar-00605-met-k-2013-File004_update.pdf. Details on HLA alleles and resolutions can be found at https://www.ebi.ac.uk/ipd/imgt/hla/ and http://hla.alleles.org/

## Code availability

The code, typing results and reference dataset used in this study is available at https://github.com/nikolasthuesen/hla-typing-benchmark.

## Supplementary figures

**Fig. S1.**
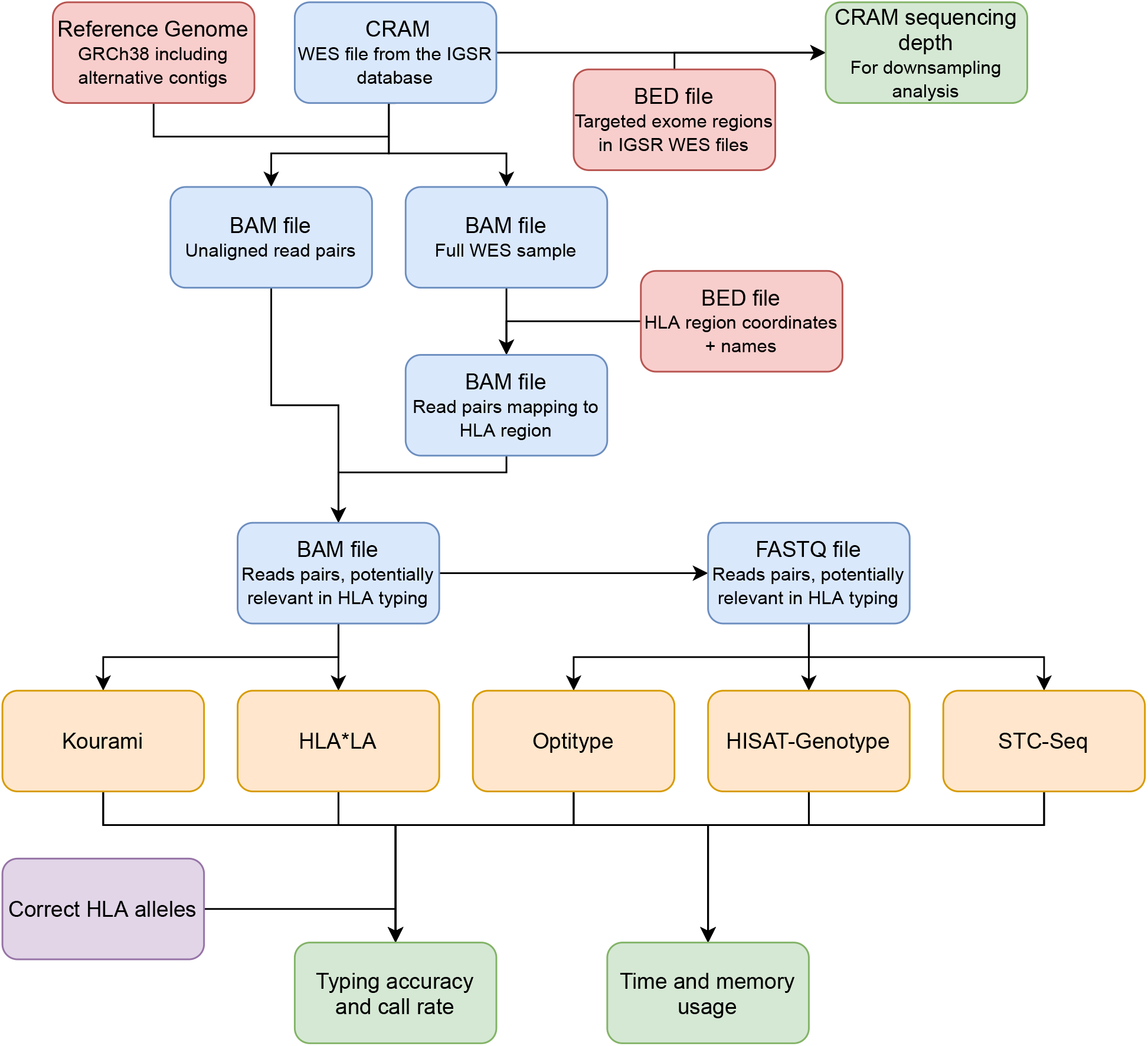
An overview of the benchmarking study. The input CRAM file from the IGSR database (gold standard WES samples) and files deriving from it are shown in blue. These are specific to the individual samples. Reference files which are identical for each sample are shown in red, the HLA typing tools in yellow, the correct HLA alleles for each sample is purple and the output files are all shown in green. For each sample (CRAM file), the sequencing depth is found (used in the downsampling analysis) and the HLA typing tools predict the HLA alleles. Predictions are compared to the correct alleles in the 1000G dataset and the computational resources used are noted for each tool. In the downsampling analysis, the found sequencing depth for each CRAM file is used, but it is not the raw CRAM file that is downsampled. Instead, the reduced BAM file (with read pairs potentially relevant in HLA typing) is downsampled, thereby avoiding the unnecessary extraction of HLA reads in the downsampling analysis. When calculating the use of memory and time, only the step specific to each tool is evaluated. This means that the performance analysis did not include the initial preprocessing measures, the extraction of HLA reads and the conversion of CRAM files to BAM files and further to FASTQ files. Figure created using https://www.diagrams.net/

**Fig. S2.**
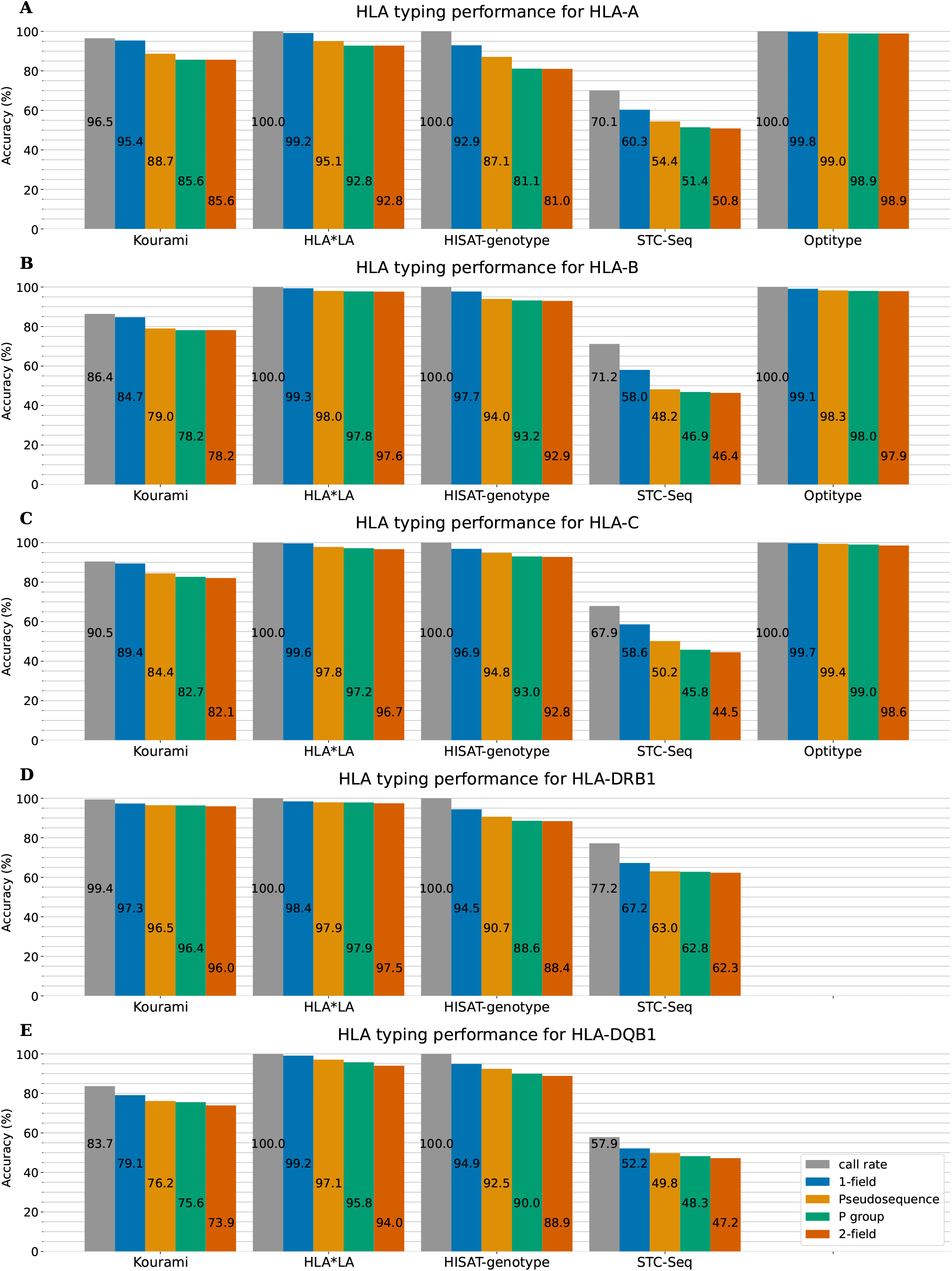
The five HLA typing tools’ typing accuracy and call rate in 1-field, pseudo-sequence, P group and 2-field resolution for HLA-A, -B, -C, -DRB1 and -DQB1.

**Fig. S3.**
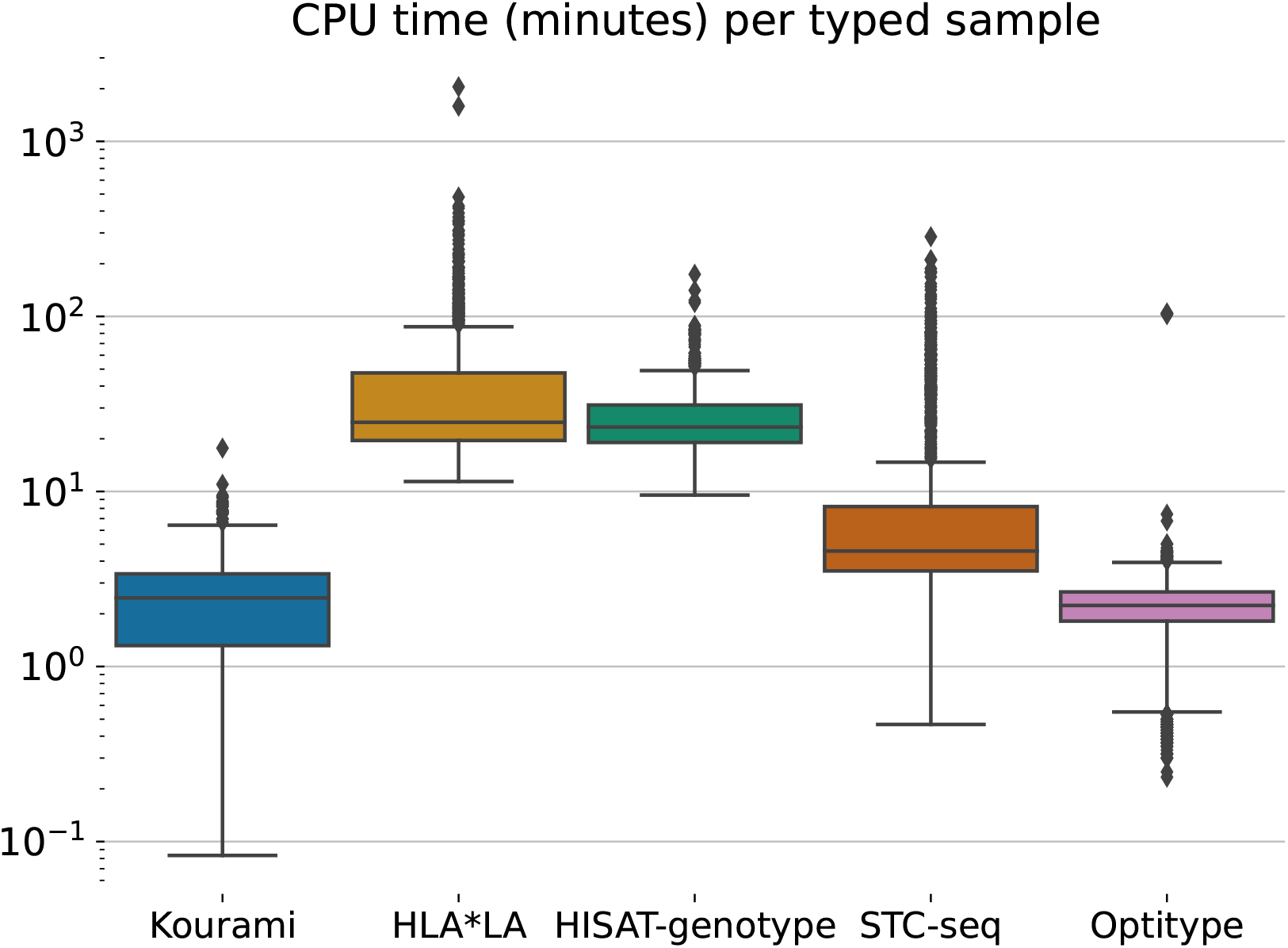
Distribution of CPU time usage for HLA typing for the 829 samples in the 1000G dataset at full coverage. HLA*LA and HISAT-genotype have the highest median CPU time usage (between 20 and 30 CPU minutes), while the rest of the tools have a median less than 10 CPU minutes. For HLA*LA, the CPU time varies a lot between samples. Some samples are typed using a similar amount of resources as is used by Kourami and Optitype, while others require over 100 times more CPU time.

**Fig. S4.**
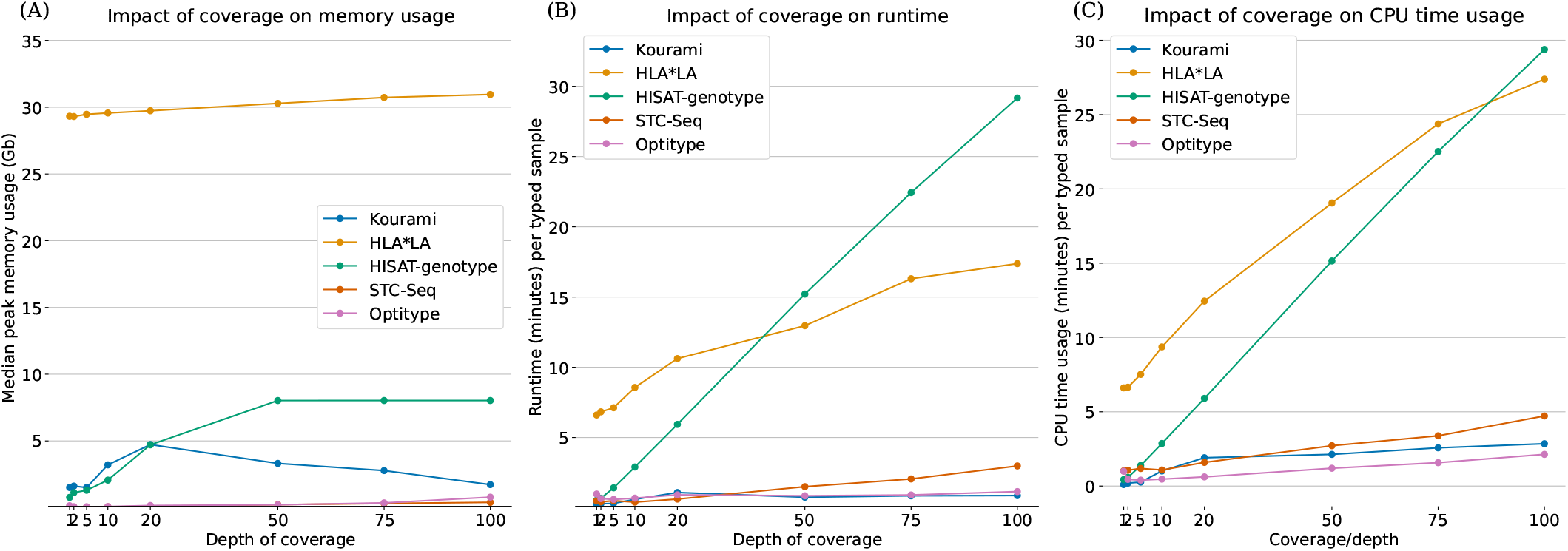
**(A):** An increased coverage generally increases the peak memory usage of the HLA typing tools, but not greatly. For HLA*LA there is almost no difference in peak memory usage for 1X and 100X. HISAT-genotype uses more memory when the coverage is increased, but only up until 8 GB. **(B) and (C):** An increase in coverage also means that it takes more time for the tools to perform HLA typing. By extrapolating the trend for HISAT-genotype, the tool would need more than an hour in real time to type samples with a coverage of above 200X (using a setup identical to the one in this benchmarking study). Keep in mind, that these times are solely for the typing step and not the extraction of HLA reads, as illustrated in figure S1.

**Fig. S5.**
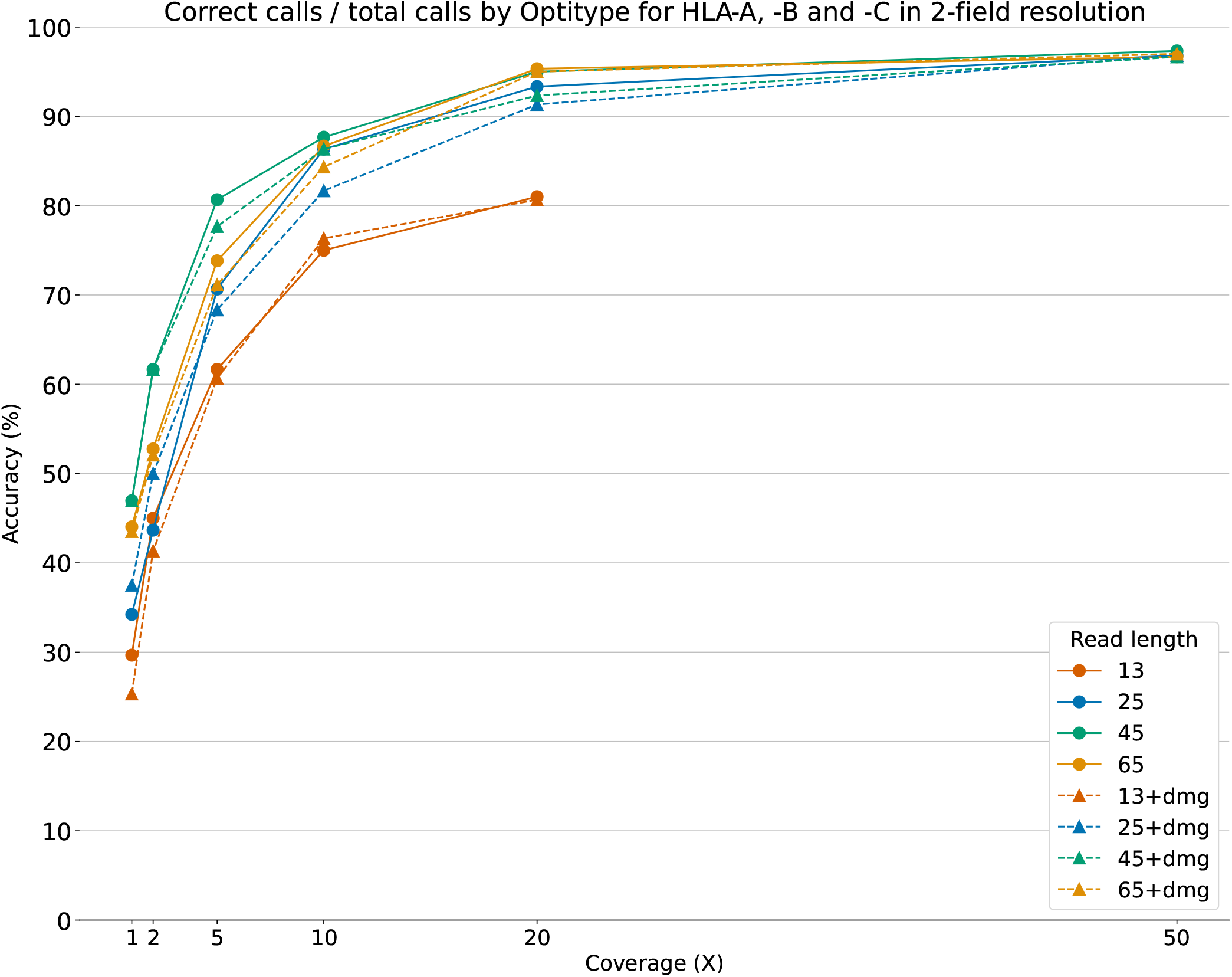
Optitype’s call rate was not 100% for all combinations of read length, coverage and with/without read damage. This is either due to the fact that Optitype did not return a call *or* a sample did not have enough data to e.g. achieve a coverage of 50X, when the read length of each read was 13. This figure shows the amount of correct calls not out of the total calls but out of the calls that Optitype did make. The trend is very similar to the one shown in figure 5, but this figure shows how confident Optitype is on a call, when it *is* made.

**Fig. S6.**
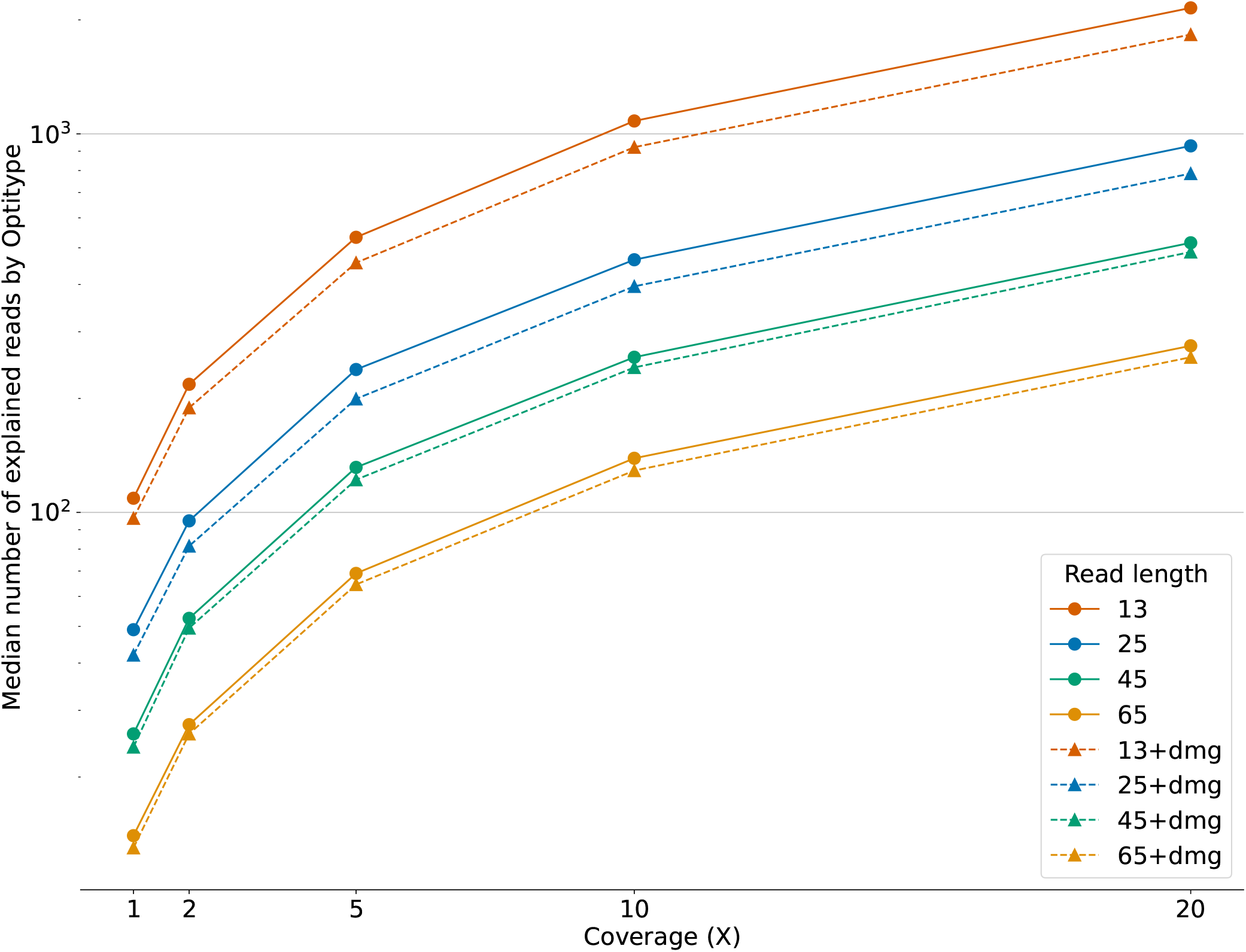
Optitype outputs the number of reads explained by its HLA allele prediction. The number of explained reads naturally depends on the coverage of the input sample. For a specific depth of coverage, there are more reads, if the read length is lower. This means, that there only are a few relevant HLA reads for samples samples with a read length of 65 and a coverage of 1X.

**Fig. S7.**
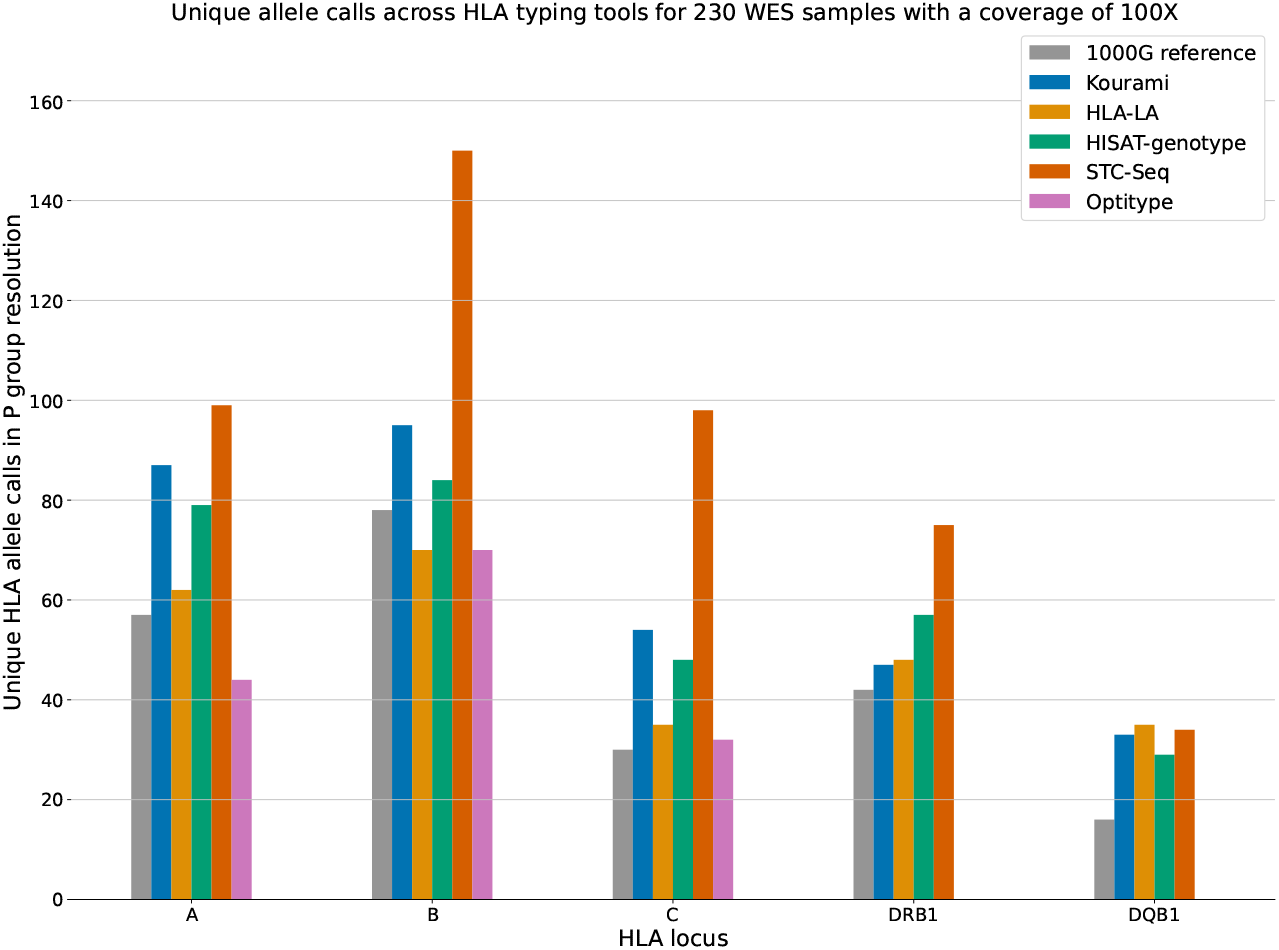
Unique HLA allele calls (in P group resolution) by the five HLA typing tools for the 230 individuals in the downsampling study, when the coverage of the samples was 100X. The true HLA alleles for these samples, as found in the 1000G dataset, is shown as well. Results are split across loci, which means that there are 460 allele calls per tool per locus.

## Footnotes

1 http://hla.alleles.org/

2 https://github.com/FRED-2/OptiType

3 https://github.com/Kingsford-Group/kourami

4 https://github.com/DiltheyLab/HLA-LA

5 https://daehwankimlab.github.io/hisat-genotype/manual/

6 https://bigd.big.ac.cn/biocode/tools/BT007068

7 http://hla.alleles.org/alleles/deleted.html

8 https://genomeinformatics.github.io/HLA-PRG-LA/

